# Megan Server: facilitating interactive access to metagenomic data on a server

**DOI:** 10.1101/2022.12.05.518498

**Authors:** Anupam Gautam, Wenhuan Zeng, Daniel H. Huson

## Abstract

**Motivation:** Metagenomic projects of large sequencing datasets (totaling hundreds of gigabytes of data). Thus, computational preprocessing and analysis are usually performed on a server rather than on a personal computer. The results of such analyses are then usually explored interactively. One approach is to use MEGAN, an interactive program that allows analysis and comparison of metagenomic datasets. Previous releases of MEGAN have required that the user first download the computed data from the server, an increasingly time-consuming process. Here we present Megan Server, a stand-alone program that serves MEGAN files to the web, using a RESTful API, facilitating interactive analysis without downloading the complete data. We describe a number of different application scenarios.

**Availability:** Megan Server is provided as a standalone program tools/megan-server in the Megan6 software suite, available at https://software-ab.cs.uni-tuebingen.de/download/megan6.

## MEGAN

Metagenomics is the study of microbiomes using DNA sequencing (1). As the throughput and cost-efficiency of sequencing technologies continue to decrease, the number and size of metagenomic samples collected in a project continues to increase.

The first three basic computations performed on such data (after quality control), are taxonomic analysis, functional analysis and comparative analysis. One major approach is based on sequence alignment (2, 3), which is computationally intensive and thus usually performed on a server or “in the cloud”.

Here we focus on the analysis of metagenomic data using the DIAMOND+MEGAN pipeline (4). This first employs the high-throughput alignment tool DIAMOND (5) to align reads against a protein reference database such as NCBI-nr (6) or AnnoTree (7, 8). This gives rise to alignments in DAA (DIAMOND alignment archive) format. The second step is to perform taxonomic and functional binning of reads using MEGAN or the associated command-line tool `daa-meganizer`. This process extends the content of the DAA files to include classification information, resulting in “meganized” DAA files.

MEGAN (“metagenome analyzer”) is an interactive application that runs on a personal computer. It comes with a number of command-line tools for analyzing data on a server. Meganized DAA files can be interactively explored and analyzed in MEGAN, and multiple files can be opened together in a single “comparison document”. MEGAN provides numerous features for interactive exploration, such as hierarchical displays of the NCBI and GTDB taxonomies and functional classifications, several types of charts, clustering algorithms, and dialogs for accessing individual reads and alignments.

### Megan Server

One drawback of MEGAN has been that files must be located on the local machine to be opened. This required that users download files from the server to their personal computer. This is becoming increasing challenging, as the size of metagenomic projects continues to grow, and it discourages collaborators from “quickly taking a look” at the data. To address this, here we present Megan Server, a light-weight, stand-alone web server that serves meganized-DAA files, and other MEGAN-associated files, to the web, using a RESTful API. During setup of the server, a root directory is specified and then all appropriate files found in or below the root directory are served. The API provides endpoints for obtaining file-related information, classification-related information, for accessing reads and matches and for administrating the server.

The service can be accessed using a web browser. For example, the URL http://maira.cs.uni-tuebingen. de:8001/megan6server/help will display a help window. However, we have implemented client code in MEGAN and the program allows the user to interactively connect to a Megan Server instance (see Figure 1). MEGAN displays a structured list of all files accessible from the server. Such files can be interactively opened and analyzes as if they were present on the local machine. While other metagenomic resources also provide APIs, they appear to lack an equally rich client for interactive exploration.

**Fig. 1.**
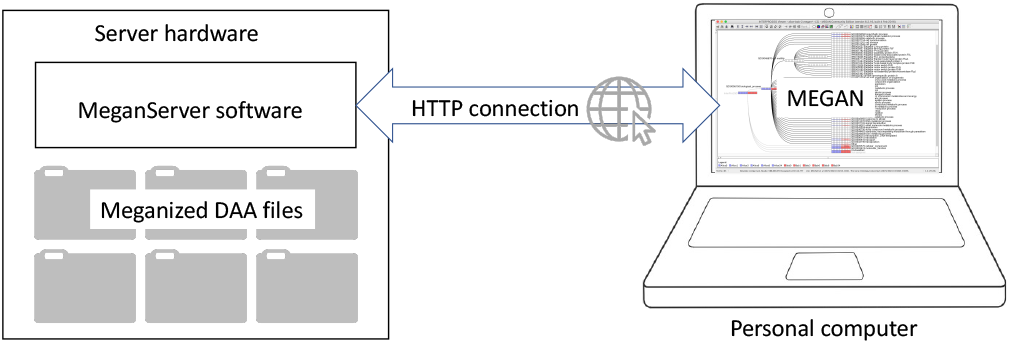
The Megan Server program runs on a server and makes meganized DAA files accessible to MEGAN running on a personal computer via the web.

Megan Server is provided in the tools directory of the MEGAN installation. To launch the server, execute the following command:

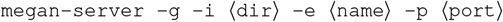

Here, the option `-g` turns on guest access, ⟨dir⟩ is the directory containing all files to be served (including all directories below), name is the endpoint ⟨name⟩, e.g. `megan6server` and ⟨port⟩ is the number of the port to be used, e.g. `8001`. See the Supplementary Material for more details. Run the program with option `-h` to see the full list of options. This program will continue to run in the background until terminated.

### Applications

Here we illustrate the use of Megan Server using four examples.

### Public Megan Server instance

We maintain a public instance of Megan Server reachable at http://maira.cs.uni-tuebingen.de:8001/megan6server/help, accessible with user-id `guest` and password `guest`. We use this instance for teaching purposes. It hosts a number of short-read datasets, including a number of kitten gut microbiomes, and over 80 long-read metagenomic datasets using data from several recently published papers.

### International collaboration using Megan Server

Together with colleagues in Brazil, we performed short-read metagenomic analysis of six fruit-waste samples to study their potential bio-surfactant bio-synthesis (9). Metagenomic analysis was performed on a server hosted in Tübingen, Germany, and were accessed from Sorocaba, Brazil, using Megan Server. It takes around 75 second to open all six samples in MEGAN via Megan Server.

### Megan Server in a cloud environment

Computational analysis of metagenomic data is often performed “in the cloud”. The “de.NBI Cloud” (German Network for Bioinformatics Infrastructure) provides configurable virtual machines (VM) for individual projects, which are accessed using a key. In one of our projects on this platform, we used a VM to run the DIAMOND+MEGAN pipeline on 172 short read samples. To access to results, we setup an instance of Megan Server on the VM. In the following, ⟨dir⟩ represents the path to the directory to be served. Access to the service requires ssh tunneling using an access key, ⟨key⟩, user name, ⟨user⟩ and the IP address of the VM, ⟨IP⟩, as follows:

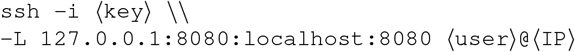

Then Megan Server is launched as follows, here name is the desired endpoint name:

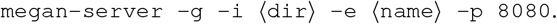

Opening all 172 samples simultaneously in MEGAN took around 10 minutes.

### Very large projects

In a current project, we are analyzing 687 short read metagenomic datasets, consisting of 4.4 TB of data in total. This data is hosted on a local linux server and we use Megan Server to access it. Loading all 687 samples into MEGAN on a laptop via Megan Server took under two hours. This loaded all classifications for all samples and most interactive analyses do not require additional contact with the server, unless the user wants to view alignments or download reads.

## Discussion

Metagenome datasets are usually analyzed on servers and are accessed via web-interfaces or downloaded and analyzed using scripts or interactive programs. Megan Server allows researchers to analyze large datasets using the interactive program MEGAN, without the need to download them.

## ACKNOWLEDGEMENTS

We acknowledge hardware support by the High Performance and Cloud Computing Group at the Zentrum für Datenverarbeitung of the University of Tübingen, the state of Baden-Württemberg through bwHPC, the German Research Foundation (DFG) through grant no. INST 37/935-1 FUGG. We also acknowledge support of the BMBF-funded de.NBI Cloud within the German Network for Bioinformatics Infrastructure (de.NBI) (031A532B, 031A533A, 031A533B, 031A534A, 031A535A, 031A537A, 031A537B, 031A537C, 031A537D, 031A538A).

## Supplementary material

Here we present an example on how to run Megan Server using data from our recently published work on fruit-waste fermentation metagenomic samples. In this study, Megan Server was used to provide access to the data for our collaborators in Brazil.

### 1: Starting a Megan Server instance

The directory to be served,, contains six DAA files (Supplementary Figure 2), which have already been processed by the DIAMOND+MEGAN pipeline. They represent fruit-waste fermentation samples ((9)). The directory is hosted on an in-house server named and MEGAN was installed on the server.

An instance of Megan Server is started by typing the following command:

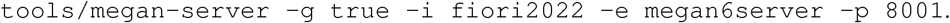

Here, the option `-g true` turns on “guest access” with user-id “guest” and password “guest”. The directory to be served (option `-i`) is `fiori2022` and the endpoint name (option `-e`) is set to `megan6server`. The option `-i 8001` sets the port to be used for serving.

As shown in Figure 2a, the software lists the address at which the server can be reached, in this example, the URL is http://osa3:8001/megan6server. Please do not attempt to access this instance of Megan Server, as it is used here only for illustration purposes. A public instance of Megan Server can be accessed here: http://maira.cs.uni-tuebingen.de:8001/megan6server/help.

**Fig. 2.**
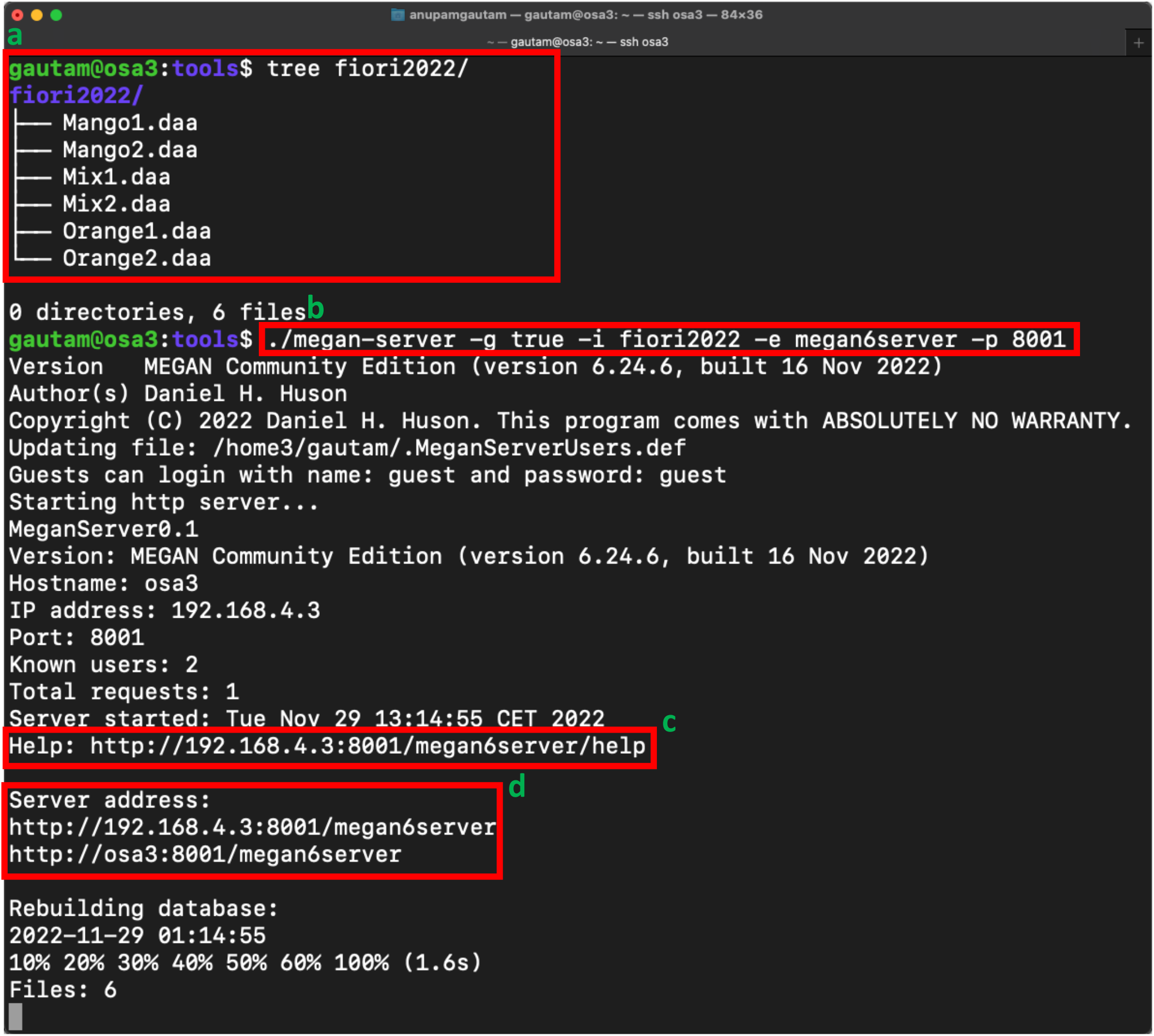
Starting a Megan Server instance on server. (a) In this example, an the directory to be served is called `fiori2022` and is located on a computer called `osa3`. (b) This command launches Megan Server and the program will serve the files to the internet via the specified port. (c) Use URL into a web browser to obtain a help screen. (d) This base URL can be used in MEGAN to contact the server.

Note that when you run the software for the first time, then you will be prompted to provide a password for the username `admin`, and the program will generate a `.MeganServerUser.def` file in your home directory, which is used to maintain user data.

### 2: Accessing the served data from your personal computer

#### 2.1: Web browser access

When Megan Server is launched, it lists an address (Figure 2c) that can be used to access the service from a web browser (Figure 3a), in this example, it is http://osa3:8001/megan6server. (This is an example URL, it is not externally reachable.) Specifying the resource `help` will display a help window (Supplementary Figure 3a). You can specify other resources listed on the help page to interact with your data. For example, specify the resource `list`, like this http://osa3:8001/megan6server/list, to obtain a list of all samples present in the served directory. The first time you access a resource other than `help`, you will be prompted for a user-id and password, such as `guest` and `guest` (Supplementary Figure 3b). In this example, we will obtain a list of the six files contained in the `fiori2022` directory (Supplementary Figure 3c).

**Fig. 3.**
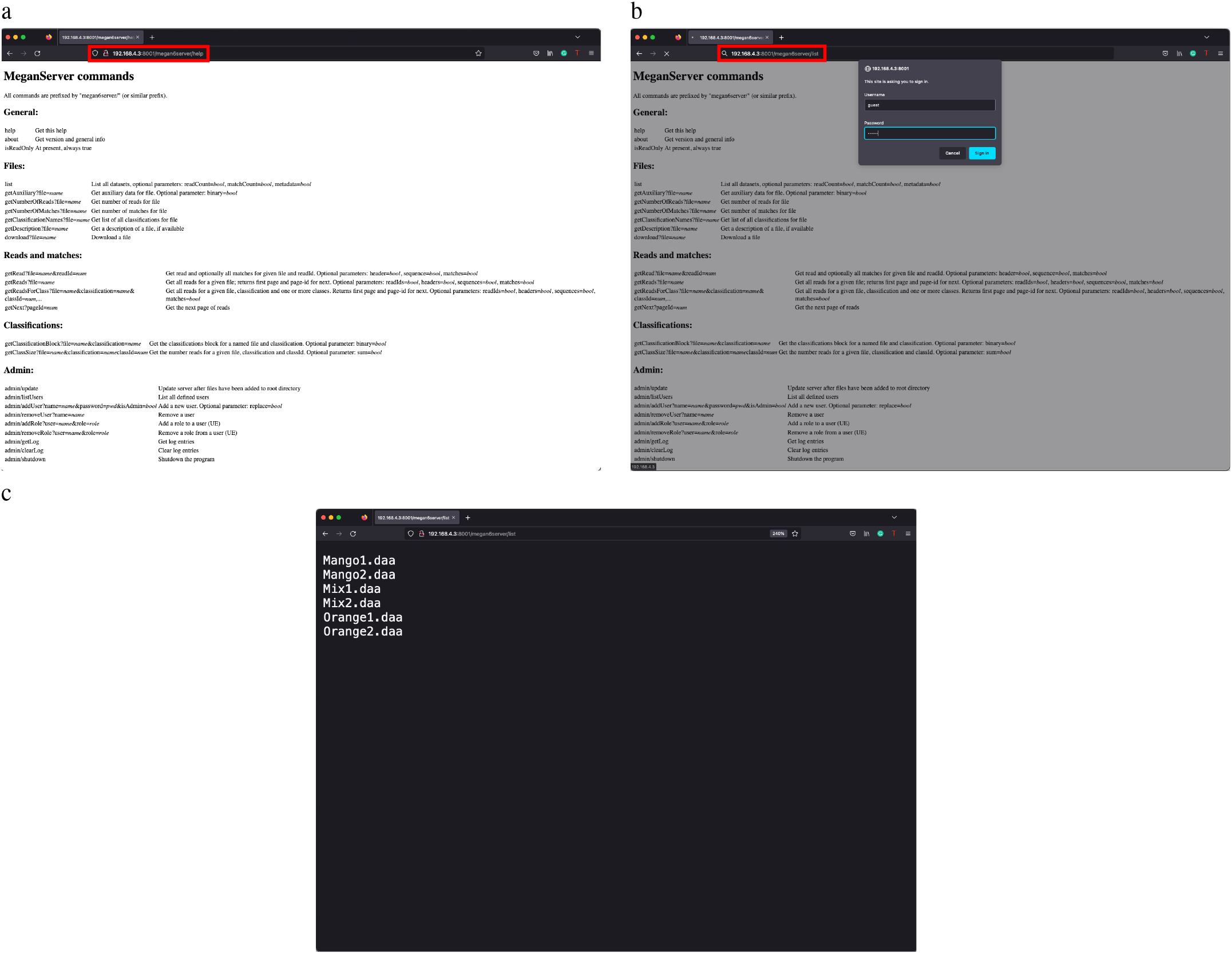
Accessing and interacting with data through the web. (a) Enter the Megan Server URL and a resource request into a web browser. For example, http://osa3:8001/megan6server/help requests the resource `help`, which results in the displaying of a help text. (b) Requesting the `list` resource, by typing http://osa3:8001/megan6server/list, results in a list of all served files (c). This may trigger an `authentication` dialog.

#### 2.2: MEGAN access to served files

Files can also be accessed using the program MEGAN, which acts as a client for the server. To contact an instance of Megan Server, select the `File` → `Open From Server…` menu item (Figure 4a). In the dialog window (Figure 4b), enter the server endpoint and user credentials (for example, user `guest` and password `guest`). The program then displays a list of all available remote files (Figure 4c). Select and open any files of interest. Multiple files can also be opened together in a single document using the `Compare` button (Supplementary Figure 4d). Then, clicking the `Apply` button will show the selected file or files in the default NCBI taxonomy viewer (Supplementary Figure 4e).

**Fig. 4.**
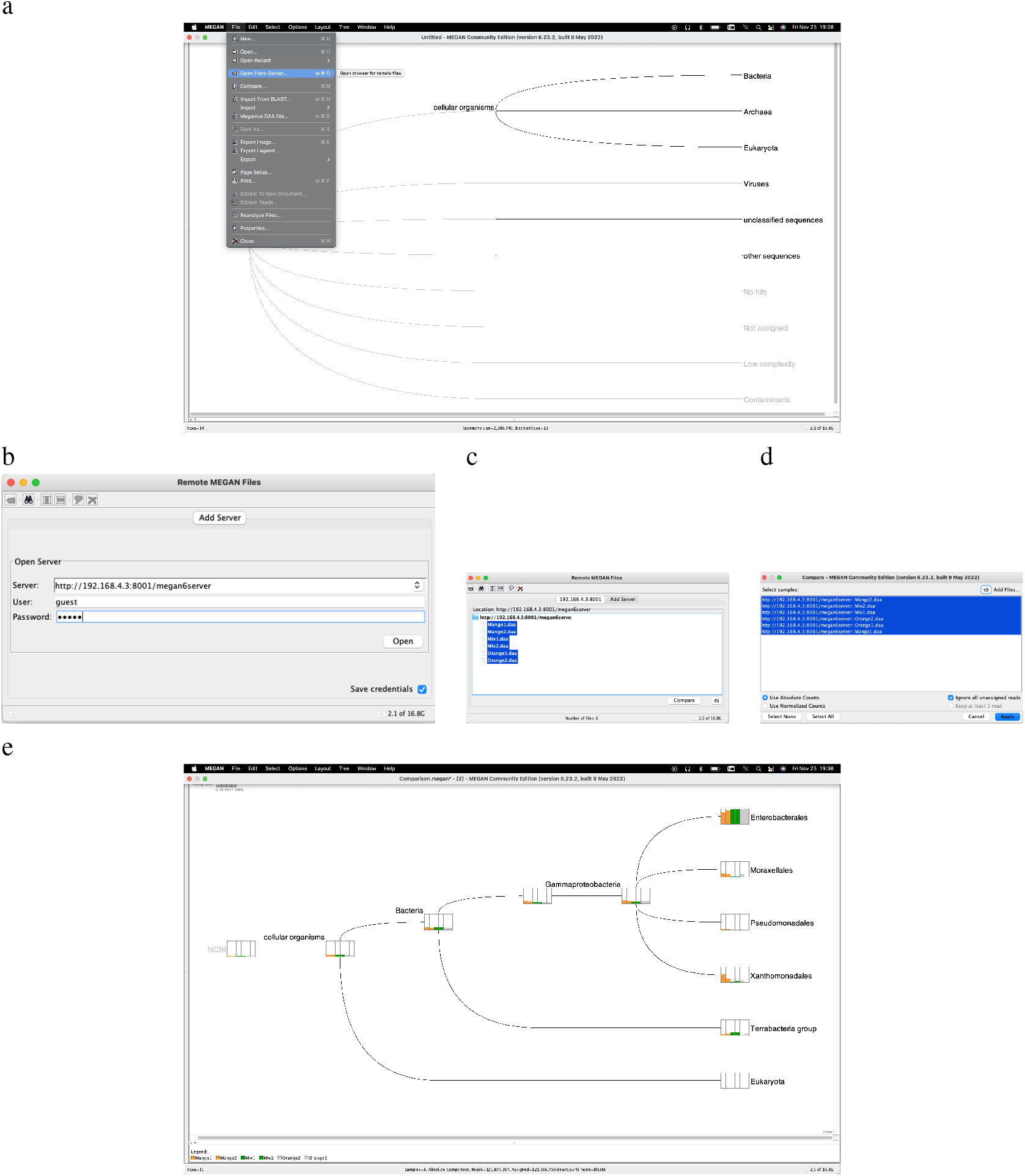
Accessing and interacting via MEGAN-GUI. (a) To access an instance of Megan Server, select the `File` *→* `Open From Server…` menu item and then (b) enter the server endpoint and user credentials. (c) After successfully contacting the service, an overview of all available remote files is displayed. (d) If more than one file is selected, then one can use the `Compare` dialog to combine multiple files into a single document. (e) Pressing `Apply` will open the files in the default NCBI taxonomy viewer.

